# Amikacin potentiator activity of zinc complexed to a pyrithione derivative with enhanced solubility

**DOI:** 10.1101/2021.10.08.463674

**Authors:** Jesus Magallon, Peter Vu, Craig Reeves, Stella Kwan, Kimberly Phan, Crista L. Oakley-Havens, Kenneth Rocha, Veronica Jimenez, María Soledad Ramirez, Marcelo E. Tolmasky

## Abstract

Resistance to amikacin in Gram-negatives is usually mediated by the 6′-*N*-acetyltransferase type Ib [AAC(6′)-Ib], which catalyzes the transfer of an acetyl group from acetyl CoA to the 6′ position of the antibiotic molecule. A path to continue the effective use of amikacin against resistant infections is to combine it with inhibitors of the inactivating reaction. We have recently observed that addition of Zn^2+^ to in-vitro enzymatic reactions, obliterates acetylation of the acceptor antibiotic. Furthermore, when added to amikacin-containing culture medium in complex to ionophores such as pyrithione (ZnPT), it prevents the growth of resistant strains. An undesired property of ZnPT is its poor water-solubility, a problem that currently affects a large percentage of newly designed drugs. Water-solubility helps drugs to dissolve in body fluids and be transported to the target location. We tested a pyrithione derivative described previously (Magda et al. Cancer Res. 2008, 68:5318-5325) that contains the amphoteric group di(ethyleneglycol)-methyl ether at position 5 (compound 5002), a modification that enhances the solubility. Compound 5002 in complex with zinc (Zn5002) was tested to assess growth inhibition of amikacin-resistant *Acinetobacter baumannii* and *Klebsiella pneumoniae* strains in the presence of the antibiotic. Zn5002 complexes in combination with amikacin at different concentrations completely inhibited growth of the tested strains. However, the concentrations needed to achieve growth inhibition were higher than those required to achieve the same results using ZnPT. Time-kill assays showed that the effect of the combination amikacin/Zn5002 was bactericidal. These results indicate that derivatives of pyrithione with enhanced water-solubility, a property that would make them drugs with better bioavailability and absorption, are a viable option for designing inhibitors of the resistance to amikacin mediated by AAC(6′)-Ib, an enzyme commonly found in the clinics.

## Introduction

Water-solubility helps drugs dissolve in body fluids and be transported to the target location ^1^. Unfortunately, about half of the chemical compounds identified as potential new medicines are poorly soluble in water ^2,3^. Currently, many efforts and techniques focus on enhancing the water-solubility of lead compounds, which illustrates this property’s importance for pharmacological tools ^1-7^.

Studies to isolate inhibitors of aminoglycoside-modifying enzymes, in particular the aminoglycoside 6′-*N*-acetyltransferase type Ib [AAC(6′)-Ib], a widely distributed enzyme that specifies resistance to the semisynthetic amikacin ^8-10^, showed that Zn^2+^ complexed to pyrithione (ZnPT) (Fig. 1) counter the action of AAC(6′)-Ib in bacterial cells in culture ^11,12^. Consequently, combinations amikacin/ZnPT produced a substantial reduction in the minimal inhibitory concentration of amikacin of AAC(6′)-Ib-containing *Acinetobacter baumannii, Escherichia coli, Enterobacter cloacae*, and *Klebsiella pneumoniae* isolates ^11-14^. However, complexes formed between pyrithione and divalent metal cations, which occur through the oxygen and sulfur atoms, have very low solubility in aqueous solvents, impairing bioavailability ^15^. To deal with this limitation, Magda et al. designed pyrithione derivatives with substitutions at position 5 to enhance their solubility in aqueous solvents ^7^. Here, we show that a complex formed between a water-soluble pyrithione derivative, compound 5002, and Zn^2+^ (Zn5002) (Fig. 1) exhibits amikacin resistance inhibitory properties similar, albeit not as robust, to those observed when testing ZnPT.

**Figure 1.**
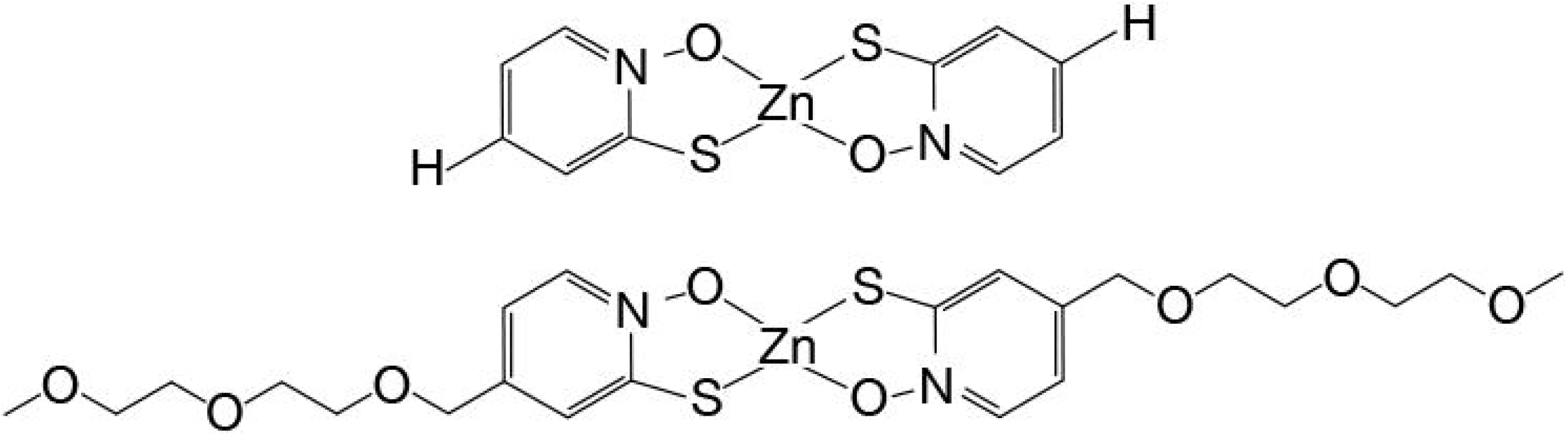
Zn5002 and ZnPT complexes chemical structures. Chemical structures of ZnPT (upper) and Zn5002 (lower).

## Results

The addition of Zn^2+^ to reaction mixtures containing AAC(6′)-Ib and aminoglycosides known to be substrates for this enzyme exerts a strong inhibition effect of the antibiotic’s acetylation ^12^. However, inhibition of resistance in cells in culture requires very high concentrations of zinc salts in the culture medium. The concentrations of zinc ions required to reverse resistance can be drastically reduced supplementing the growth medium with the complex ZnPT ^11,12^. An inconvenience to develop ZnPT as an adjuvant to aminoglycosides to treat resistant bacteria is its poor solubility in water. Previous work by Magda et al. showed that substituting the hydrogen at position 5 of pyrithione by some amphoteric chemical groups results in derivatives with higher water-solubility that can still diffuse across the membrane ^7^. We synthesized compound 5002, in which the hydrogen at position 5 is replaced by di(ethyleneglycol)-methyl ether group (Fig. 1). This compound was complexed to Zn^2+^ (Zn5002) and tested as a potentiator to amikacin to overcome resistance in AAC(6′)-Ib-carrying *A. baumannii*, and *K. pneumoniae* cells.

All four strains tested were cultured in the presence of amikacin, the ionophore-zinc complex, or a combination of both compounds at different concentrations. Fig. 2 shows the growth curves corresponding to the combinations that include the minimum possible concentration of each component to inhibit growth completely. The figure also shows that when none or only one of the components was used to supplement the Mueller-Hinton broth there was healthy bacterial growth. Although the concentrations required to inhibit growth vary from strain to strain, there was an appropriate combination in all cases such that the individual components did not impede growth. It can also be noted that the ZnPT concentration necessary to overcome the resistance to amikacin is consistently lower than that of Zn5002.

**Figure 2.**
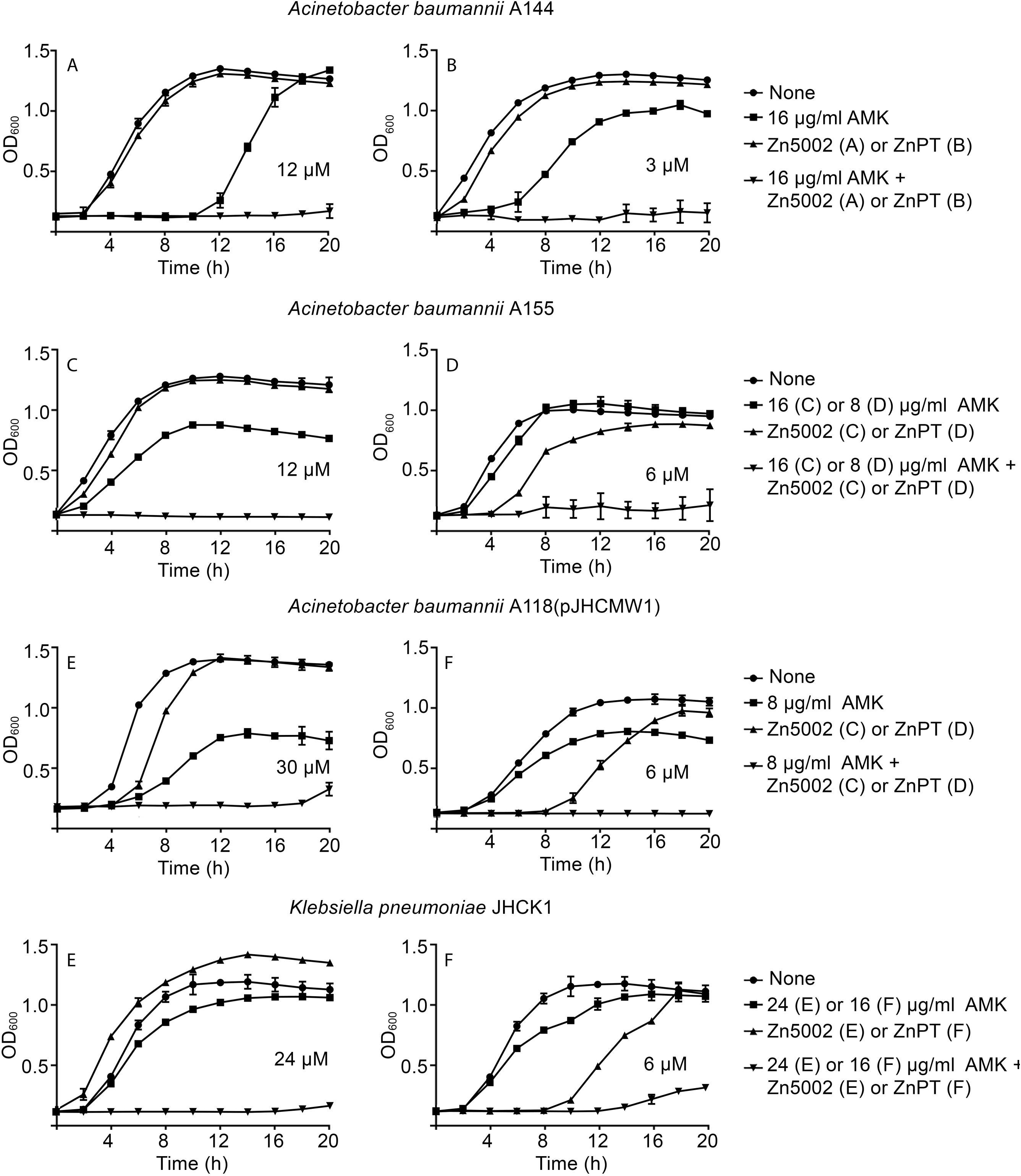
Effect of Zn5002 and ZnPT complexes on resistance to amikacin. *A. baumannii* A155, A144, A118(pJHCMW1), *K. pneumoniae* JHCK1, or *E. coli* TOP10(pNW1) were cultured in 100 μl Mueller-Hinton broth in microtiter plates at 37°C with the additions indicated to the right of each panel, and the OD_600_ was periodically measured. All cultures contained 0.5 % DMSO. AMK, amikacin.

The results obtained in the experiments described above indicate that the complex Zn5002, as we showed before for ZnPT, is responsible for the phenotypic conversion to amikacin susceptibility in bacterial pathogens harboring the resistance enzyme AAC(6′)- Ib. However, these experiments did not inform about the bactericidal or bacteriostatic effect of the combination. Therefore, we carried out time-kill assays to confirm that the inhibition of growth observed in the presence of Zn5002 and the antibiotic is due to a bactericidal effect. For comparison, we carried out another series of assays using amikacin and ZnPT. Fig. 3 shows that the addition of amikacin and Zn5002 or ZnPT is followed by rapid loss of bacterial cell viability. Conversely, addition of amikacin or zinc-ionophore alone did not result in cell death. These assays showed that amikacin has a robust bactericidal activity on the AAC(6′)-Ib-carrying *A. baumannii* A144, A155, A118(pJHCMW1), and *K. pneumoniae* JHCK1 strains when administered in combination with the complexes.

**Figure 3.**
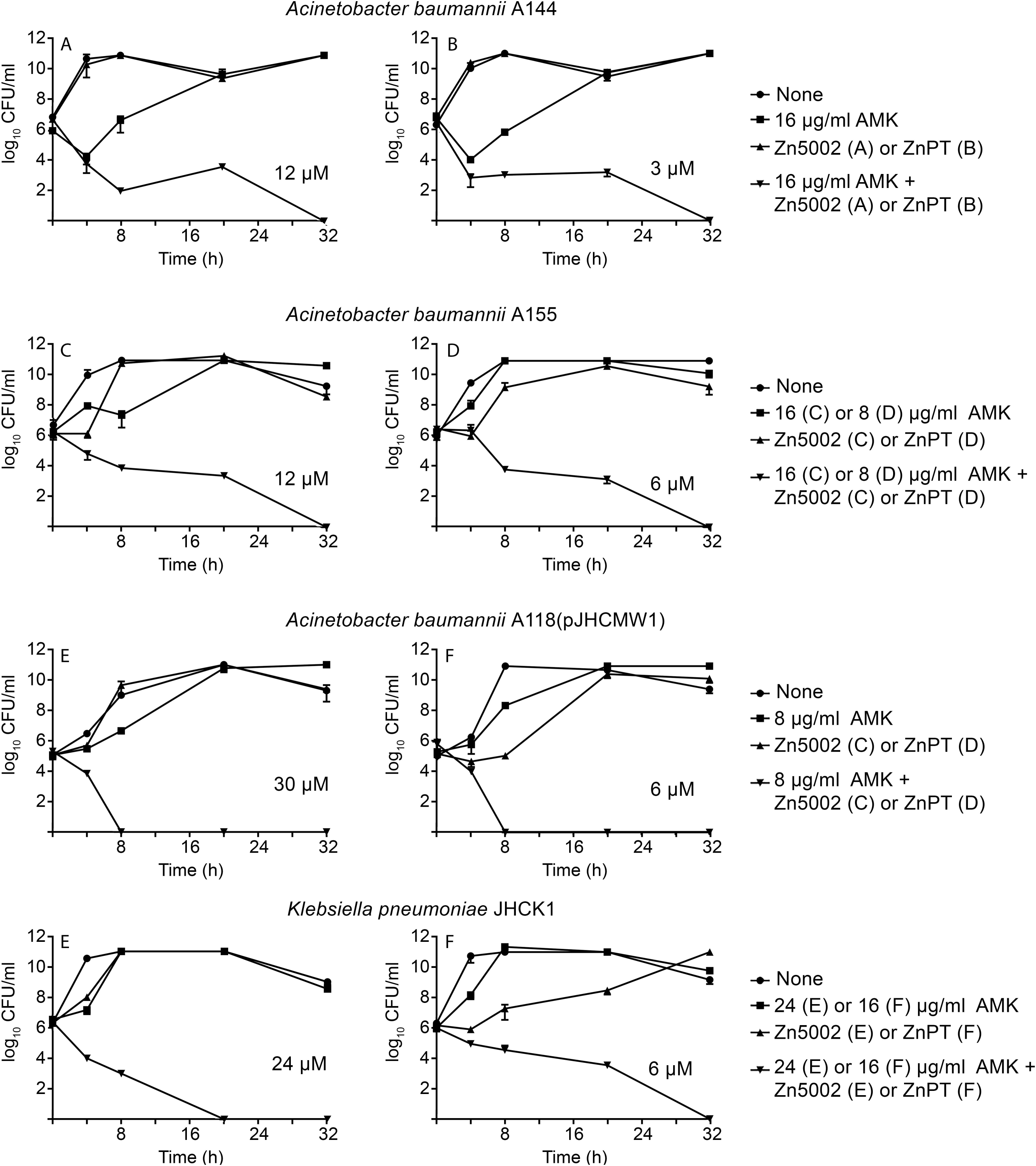
Time-kill assay curves for amikacin in the presence of Zn5002 and ZnPT. *A. baumannii* A155, A144, A118(pJHCMW1), *K. pneumoniae* JHCK1, or *E. coli* TOP10(pNW1) were cultured in 100 μl Mueller-Hinton containing 0.5% DMSO until they reach 10^6^ CFU/ml. At this moment the cultures were supplemented with the additions indicated to the right of each panel, the cultures were continued at 37°C with shaking and samples were removed periodically to determine CFU/ml. AMK, amikacin.

## Discussion

Water-solubility is a desirable characteristic of drugs for enhanced bioavailability ^2,3^. Various routes of administration, such as oral or parenteral, depend on the drug water solubility to be viable options ^5,16^. Drugs that readily dissolve in the aqueous body fluids are more efficient in reaching the desired concentrations, being transported to, and reaching their target ^1^. These characteristics make them therapeutically effective without the need to use high doses that could be the cause of secondary effects ^1^. Conversely, low water-solubility is the cause of failure of numerous drug candidates ^1^.

Amikacin is an aminoglycoside most commonly administered intravenously and intramuscularly, yet other routes are also utilized, such as intrathecal, intraventricular, topical, and inhaled ^8,17,18^. In our quest to identify compounds that inhibit the AAC(6′)-Ib amikacin-disabling action, we recently found that various cations effectively interfere with the enzymatic inactivation ^11,12,19-21^. In the case of Zn^2+^, the concentrations needed to inhibit growth of amikacin-resistant cells in the presence of the antibiotic are significantly reduced if the cation is added to the growth media in complex with ionophores ^11-13,20-22^. A very effective complex to reduce amikacin resistance levels in various bacteria is ZnPT, a compound already being researched and repurposed for cancer treatments and that has low toxicity when tested on mice ^23,24^. However, a drawback is its poor solubility in aqueous media and low bioavailability ^7,25^. Addition of an amphoteric group, di(ethyleneglycol)-methyl ether, to position 5 of pyrithione (compound 5002) enhances the chemical’s solubility in water without increasing toxicity (Fig. 1) ^7^. A comparison of the complexes Zn5002 and ZnPT showed that both compounds act as adjuvants to amikacin. The addition of Zn5002 plus amikacin to the nutrient medium inhibits growth and has a bactericidal effect. The active concentrations of the components, amikacin and Zn-ionophore complex, varied from strain to strain.

This characteristic can be due to *aac(6*′*)-Ib* gene dosage or other mechanisms or properties that may help the resistance, such as efflux pumps or low permeability. Inspection of the results indicates that the active concentrations of Zn5002 were consistently higher than those of ZnPT, suggesting that a reduction in activity accompanied the gain in solubility in aqueous solutions. However, the fact that a highly water-soluble derivative of ZnPT conserved the activity indicates that further research will permit us to design other robust adjuvants with high water-solubility. Those compounds will be strong potentiators to aminoglycosides to overcome resistance.

## Methods

### Bacterial strains

The bacterial strains used in this study were *A. baumannii* A155 ^26^, A144 ^27^, and A118(pJHCMW1) ^28^, and *K. pneumoniae* JHCK1 ^29^. *A. baumannii* A155 and A144 are multidrug-resistant and include *aac(6*′*)-Ib* in their genomes ^26,27^. *A. baumannii* A118 is a blood isolate characterized for being susceptible to most antibiotics ^28^. This strain was transformed with pJHCMW1, a plasmid that carries *aac(6*′*)-Ib* ^30^. *K. pneumoniae* JHCK1 is a multidrug-resistant isolate from cerebrospinal fluid of a neonate with meningitis ^31^.

### General procedures

Routine bacterial cultures were carried out in L broth (Lennox, 1% tryptone, 0.5% yeast extract, 0.5% NaCl), with the addition of 2% agar for plates. was tested inoculating 100-μl Mueller-Hinton broth in microtiter plates with the specified additions using the BioTek Synergy 5 microplate reader

Inhibition of growth was determined by inoculating Mueller-Hinton broth (100-μl) containing the indicated additions. The microtiter plates were incubated with shaking at 37°C in a BioTek Synergy 5 microplate reader as previously described ^20^. The cultures’ optical density values at 600 nm (OD_600_) were determined at regular intervals. ZnPT was purchased from MilliporeSigma, and Zn5002 was synthesized and purified to 97.87% by BioSynthesis Inc. All cultures to determine the action of zinc-ionophore complexes inhibition of resistance to amikacin or bactericidal effect included 0.5% dimethylsulfoxide (MilliporeSigma). Time-kill assays were performed as before ^20^. Briefly, cells were cultured in Mueller-Hinton broth until they reached 10^6^ CFU/ml. At this time, the compounds to be tested were added, and the cultures were continued at 37°C with shaking. The number of cells was determined by taking aliquots after 0, 4, 8, 20, and 32 h.

## Data availability

Bacterial strains used in this work are available upon request.

## ACKNOWLEDGEMENTS

This work was supported by Public Health Service grants 2R15AI047115 (M.E.T.) from the National Institute of Allergy and Infectious Diseases, National Institutes of Health, SC3GM125556 (M.S.R.) from the National Institute of General Medical Sciences, National Institutes of Health, and California State University Fullerton.

## COMPETING INTERESTS

The author(s) declare no competing interests.

## AUTHOR CONTRIBUTIONS

Conceptualization, M.E.T.; formal analysis, J.M., M.S.R., and M.E.T.; funding acquisition, M.S.R. and M.E.T.; methodology, J.M., P.V., C.R., S.K., K.P., C.L. O.-H., K. R., V.J., M.S.R., and M.E.T.; resources, M.S.R., and M.E.T.; writing—original draft preparation, M.E.T.; writing—review and editing, J.M., M.E.T., and M.S.R. All authors have read and agreed to the published version of the manuscript.

## Notes

### Competing Interest Statement

The authors have declared no competing interest.

